# Lipofuscin-like autofluorescence within microglia and its impact on studying microglial engulfment

**DOI:** 10.1101/2023.02.28.530224

**Authors:** Jacob M. Stillman, Francisco M. Lopes, Jing-Ping Lin, Kevin Hu, Daniel S. Reich, Dorothy P. Schafer

**Affiliations:** Department of Neurobiology, Brudnick Neuropsychiatric Research Institute, University of Massachusetts Chan Medical School, Worcester, MA 01605, USA; Department of Neurobiology, Brudnik Neuropsychiatric Research Institute, University of Massachusetts Chan Medical School, Worcester, MA, USA; University of Massachusetts Chan Morningside Graduate School of Biomedical Sciences, Neuroscience Program, Worcester, MA, USA; Translational Neuroradiology Section, National Institute of Neurological Disorders and Stroke, National Institutes of Health, Bethesda, MD 20892, USA

## Abstract

Engulfment of cellular material and proteins is a key function for microglia, a resident macrophage of the central nervous system (CNS). Among the techniques used to measure microglial engulfment, confocal light microscopy has been used the most extensively. Here, we show that autofluorescence (AF), likely due to lipofuscin and typically associated with aging, can also be detected within microglial lysosomes in the young mouse brain by light microscopy. This lipofuscin-AF signal accumulates first within microglia and increases with age, but it is not exacerbated by amyloid beta-related neurodegeneration. We further show that this lipofuscin-AF signal within microglia can confound the interpretation of antibody-labeled synaptic material within microglia in young adult mice. Finally, we implement a robust strategy to quench AF in mouse, marmoset, and human brain tissue.

## Introduction

Microglia are highly phagocytic tissue-resident macrophages of the central nervous system (CNS). While the phagocytic activity of microglia has historically been attributed to clearing dead or dying cells, the list of microglial phagocytic substrates has expanded in recent years to include synaptic material^1–4^, extracellular matrix proteins^5^, and protein aggregates (amyloid beta, tau, etc.)^6^. From this work, the engulfment of cellular and protein material by microglia has been shown to regulate synaptic connectivity and modulate neurodegenerative phenotypes^1–4^. Microglial engulfment is also an emerging target for therapeutic intervention in diseases ranging from Alzheimer’s disease to schizophrenia^7–9^. Therefore, it is critical that the analysis of microglial engulfment of cellular and protein substrates is performed with the highest rigor.

Confocal light microscopy has become a standard method to measure microglial engulfment function in tissues and cells^10,11^. A potential confound of these studies is autofluorescence (AF) in brain tissue. Likely the largest source of AF in tissues is lipofuscin. Lipofuscin is a mixture of highly oxidized lipids, misfolded proteins, and metals, which accumulates with age within lysosomal compartments^12–14^. These lipofuscin aggregates autofluoresce across the fluorescent spectrum, making it challenging to image fluorescently labeled cells and molecules by light microscopy^15,16,17,18^. In microglia, the aggregation of lipofuscin can be induced by incomplete myelin digestion and disruption of the lysosomal pathway, which implicates phagocytosis of cellular material as a key mechanism leading to lipofuscin buildup^19^. Further, lipofuscin accumulation in microglia is an age-dependent process and it has been estimated that AF-positive microglia, which is likely lipofuscin, outnumber AF-negative microglia by greater than two-fold in 6-month-old mice^17,20^. However, recently AF attributed to lipofuscin has been shown within microglia lysosomes as early as 7–9 weeks^17,18^. Thus, it is important to consider the potential confound that lipofuscin in microglia can be misinterpreted as engulfed cellular and protein material by microglia, leading to false positive detection of engulfed material within microglia.

Here, we assessed AF, which is likely from lipofuscin (lipofuscin-like AF or lipo-AF), within microglia using confocal light microscopy across the developing, adult, aged and diseased mouse cortex. Our data show that microglia are the first resident CNS cell type to accumulate lipo-AF, with a small amount of AF detected within microglia in the postnatal and juvenile mouse cortex. We also provide evidence that, if not taken into consideration, lipo-AF can potentially be misinterpreted as engulfed material within microglia, even in the young adult brain. Finally, we provide an adaptable pre-staining AF quenching protocol that preserves immunofluorescent antibody signal. This protocol can further be applied across species, including mouse, marmoset, and human brain tissue.

## Results

### Lipofuscin-like autofluorescence accumulates within microglial lysosomes with age independent of neurodegeneration

We began imaging tissue at postnatal day 90 (P90) when other studies have shown a significant accumulation of AF, likely due to lipofuscin, within microglia in the mouse brain by light microscopy^17,18,21^. We focused our imaging in the somatosensory cortex and neighboring visual and auditory cortices as this was a large region that could be easily identified across ages and is known to undergo neurodegeneration (Fig. 1). Unstained tissue (Fig. 1a-b) or tissue immunostained to label microglia (anti-IBA1) were imaged (Fig. 1d-e). The AF signal within the unstained cortex at P90 was observed with a 488 nm laser (Green, Band Pass Filter (BP) 525/50), 561nm laser (Red; BP 629/62), and 638nm laser (Far-red; BP 690/50), but not the 350 nm laser (Blue, BP 450/50) (Fig. 1b,c). Further, when assessed within anti-IBA1 immunostained tissue, this AF was largely localized to CD68+ lysosomes (Supplementary Fig. 1). Because of these excitation and emission properties of the AF signal and the localization of the AF signal within lysosomes, it is most likely due to lipofuscin^13,16,18,21–23^. Other AF molecules generally have a tighter excitation and emission spectra and are not localized specifically to lysosomal compartments^23^. However, there does not exist a highly specific stain for lipofuscin, therefore, we refer to it as lipofuscin-like AF or lipo-AF.

**Figure 1.**
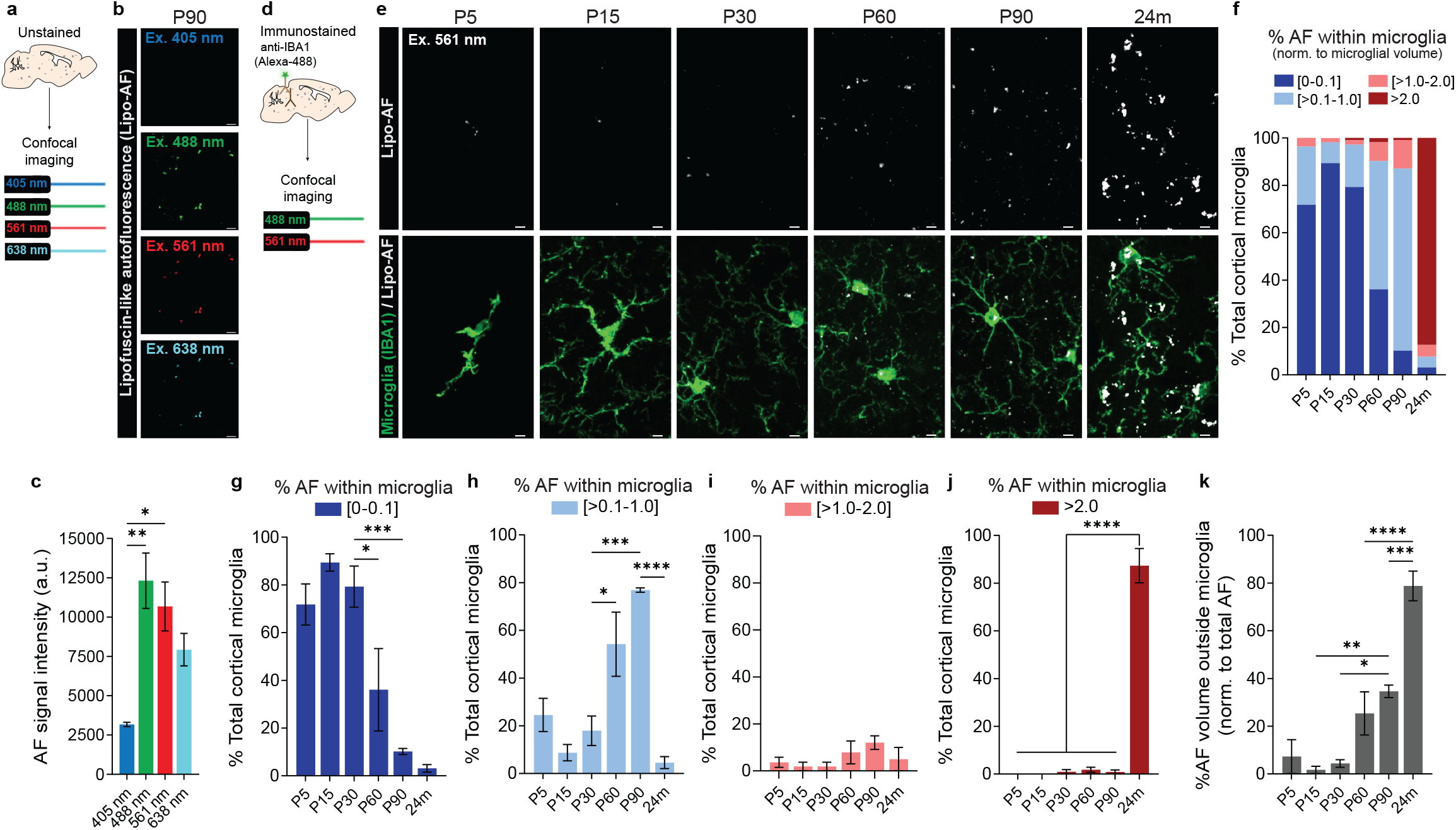
Cortical microglia accumulate lipofuscin-like autofluorescence (lipo-AF) throughout life. **a** Method for assessing lipo-AF across laser lines in unstained tissue. **b** Representative images of lipo-AF signal in unstained tissue in the young adult P90 cortex. Scale bar = 5 μm. **c** Quantification of lipo-AF signal intensity across laser lines. Data presented as mean ± SEM, n = 3 mice. One-way ANOVA with Tukey multiple comparisons test (*F* = 9.711, *df* = 11); *p<0.05 **p<0.01. **d** Method for assessing lipo-AF within microglia. **e** Representative images of anti-IBA1 immunolabelled microglia (green) and lipo-AF (white) in the developing (P5, P15), young adult (P30, P60, P90), and aged (24 months) somatosensory cortex and neighboring visual and auditory cortices. The AF was excited by the 561 nm laser line. Scale bars = 5 μm. **f** Quantification of the percentage of cortical microglial with [0-0.1]% (dark blue), [>0.1-1.0]% (light blue), [>1.0-2.0]% (pink), or >2.0% (dark red) of their total volume occupied by lipo-AF. **f** Data represented as a mean, n = 4 mice P5–P90, n = 3 mice 24m. ***g-j*** The same data as **f**, but each bin is graphed separately. Data are presented as a mean ± SEM. One-way ANOVA with Tukey multiple comparisons test (**g**, *F* = 15.49, *df* = 22; **h**, *F* = 15.53, *df* = 22; **i**, *F* = 1.584, *df* = 22; **j**, *F* = 129.9, *df* = 22); *p<0.05 ***p<0.001 ****p<0.0001. **k** Quantification of the percentage of lipo-AF volume that is outside of microglia. Data represented as mean ± SEM, n = 4 mice P5–P90, n = 3 mice 24m. One-way ANOVA with Tukey multiple comparisons test (*F* = 24.51, *df* = 22); *p<0.05 **p<0.01 ***p<0.001 ****p<0.0001.

We then extended our analyses to earlier developmental timepoints and assessed the liop-AF within anti-IBA-1 immunostained tissue (Fig. 1d-k). For simplicity, we continued our analyses using the 561nm laser (BP 629/62). From P5 to P30, most microglia had low to no detectable lipo-AF within their cytoplasm (0–0.1% of microglia volume) (Fig. 1f (dark blue) and g). However, there was a subset of microglia at these early ages (P5=24.5%±7%, P15=8.9%±3.4, P30= 17.9%±6.2) that contained more lipo-AF within their cytoplasm (Fig. 1f (light blue and pink) and h-i). By P60, there was a significant increase of lipo-AF within microglia compared to younger ages with 54.2%±13.4 of the total microglia with >0.1–1% of their volume occupied by lipo-AF (Fig. 1f (light blue) and h). This was further increased to 76.9%±0.9 of microglia with >0.1–1% of their volume occupied by lipo-AF by P90 (Fig. 1f (light blue) and 1h). As expected, most microglia (87.3%±7.2) from aged, 24-month-old brain had >2.0% of their volume occupied by lipo-AF (Fig. 1f (dark red) and j). Interestingly, lipo-AF was largely localized within microglia in the young adult brain, but by 24 months, a significantly higher percentage of lipo-AF was localized outside microglia (Fig. 1e,k).

Increased microglial engulfment of cellular material and protein aggregates has also been shown in the context of neurodegeneration, and microglial engulfment of myelin has been suggested to drive the accumulation of lipo-AF in microglia^12,19^. We, therefore, next assessed microglial AF accumulation in an Alzheimer’s disease (AD)-relevant mouse model, the 5XFAD model (Fig. 2). Surprisingly, while AF was observed in microglia in the somatosensory cortex of 9-month-old 5XFAD mice, it was comparable to 9-month wild-type (WT) controls (Fig. 2c-g).

**Figure 2.**
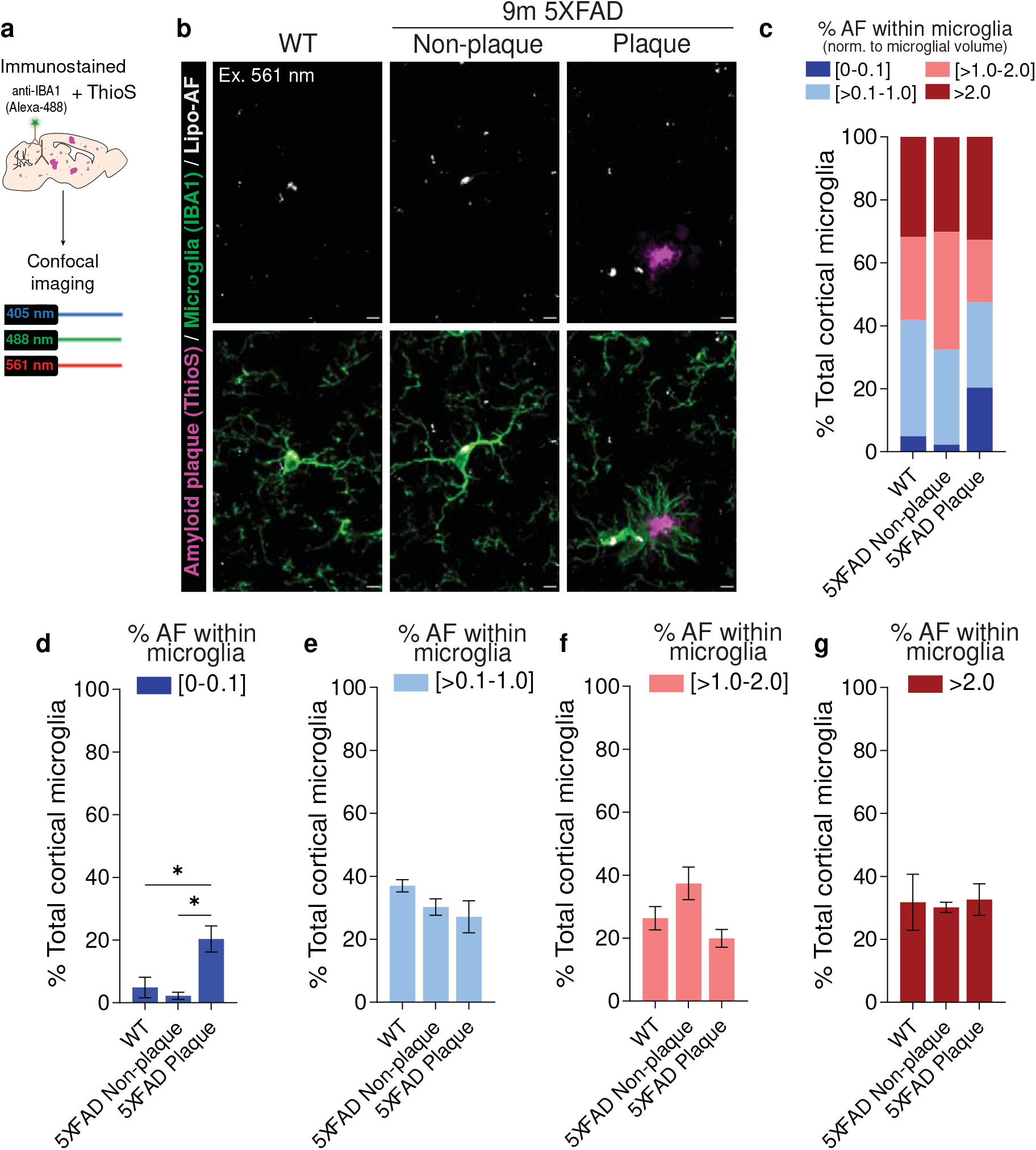
Microglial accumulation of lipo-AF signal is not exacerbated in a mouse model of neurodegeneration. **a** Methods for assessing lipo-AF in microglia in the 5XFAD mouse model. **b** Representative images of anti-IBA1 immunolabelled microglia (green), Thioflavin S labeled amyloid plaques (ThioS, magenta), and lipo-AF (white) in the somatosensory cortex of 9-month-old 5XFAD mice and wild-type (WT) littermates. The lipo-AF was excited by a 561 nm laser. Scale bars = 5 μm. **c** Quantification of the percentage of cortical microglial with [0-0.1]% (dark blue), [>0.1-1.0]% (light blue), [>1.0-2.0]% (pink), or >2.0% (dark red) of their total volume occupied by AF. Data are represented as the mean, n = 3 mice. **d-g** Same data as **c**, but each bin is graphed separately. Data are represented as mean ± SEM. One-way ANOVA with Tukey multiple comparisons test (**d**, *F* = 9.74, *df* = 8; **e**, *F* = 2.095, *df* = 8; **f**, *F* = 4.813, *df* = 8; **g**, *F* =0.04357,*df* = 8); *p<0.05.

Moreover, in Aβ plaque-enriched cortical regions, a significantly higher percentage of microglia had little to no detectable lipo-AF (0–0.1%) within their cytoplasm compared to WT mice or non-plaque-associated microglia in 5XFAD mice (Fig. 2c, dark blue and d).

Together, these data demonstrate that microglia accumulate lipo-AF prior to other cell types and earlier than previously appreciated. While lipo-AF accumulates inside and outside microglia with age, there does not appear to be a significant increase in lipo-AF accumulation within cortical microglia in the presence of Aβ plaques in 5XFAD mice.

### A reliable protocol to quench lipo-AF in mouse brain tissue

Considering the potential for lipo-AF to confound downstream analyses, we explored protocols to reduce microglial AF in brain tissue. Previous groups have used a commercially available derivative of Sudan Black to eliminate AF in tissues^18^. We repeated these experiments at P90, a timepoint when lipo-AF accumulation in microglia is significantly increased (Fig. 1). Using the commercially available reagent TrueBlack Plus^™^ after immunostaining, we then imaged with a 561nm laser (BP 629/62) and identified a significant decrease in microglial lipo-AF (Fig. 3a-f). That is, with quenching, there was a significant decrease in microglia with detectable lipo-AF (>0.1–2%) (Fig. 3c light blue, pink and e-f) and a significant increase in microglia with negligible to low lipo-AF (0–0.1%, Fig. 3c dark blue and d). However, this TrueBlack Plus^™^ quenching protocol also resulted in a significant decrease in the intensity of an immunostained protein of interest in the tissue (anti-P2RY12; Fig. 3b, g). Therefore, we took steps to improve this methodology using a commercially available MERSCOPE photobleacher device typically used for multiplexed error-robust fluorescence in situ hybridization (MERFISH). This device uses light to photobleach samples prior to immunostaining. Other groups have used LED light-based systems to achieve similar effects in tissues^21,24^. After incubating mouse brain sections in photobleaching light for 12 hours, we proceeded with our standard immunostaining protocol (Fig. 3h). In contrast to chemical quenching methods (Fig. 3a-g), this photobleaching method significantly eliminated lipo-AF signal within P90 cortical microglia (Fig. 3i-m) without compromising the fluorescent signal of anti-P2RY12+ immunostaining (Fig. 3i,n).

**Figure 3.**
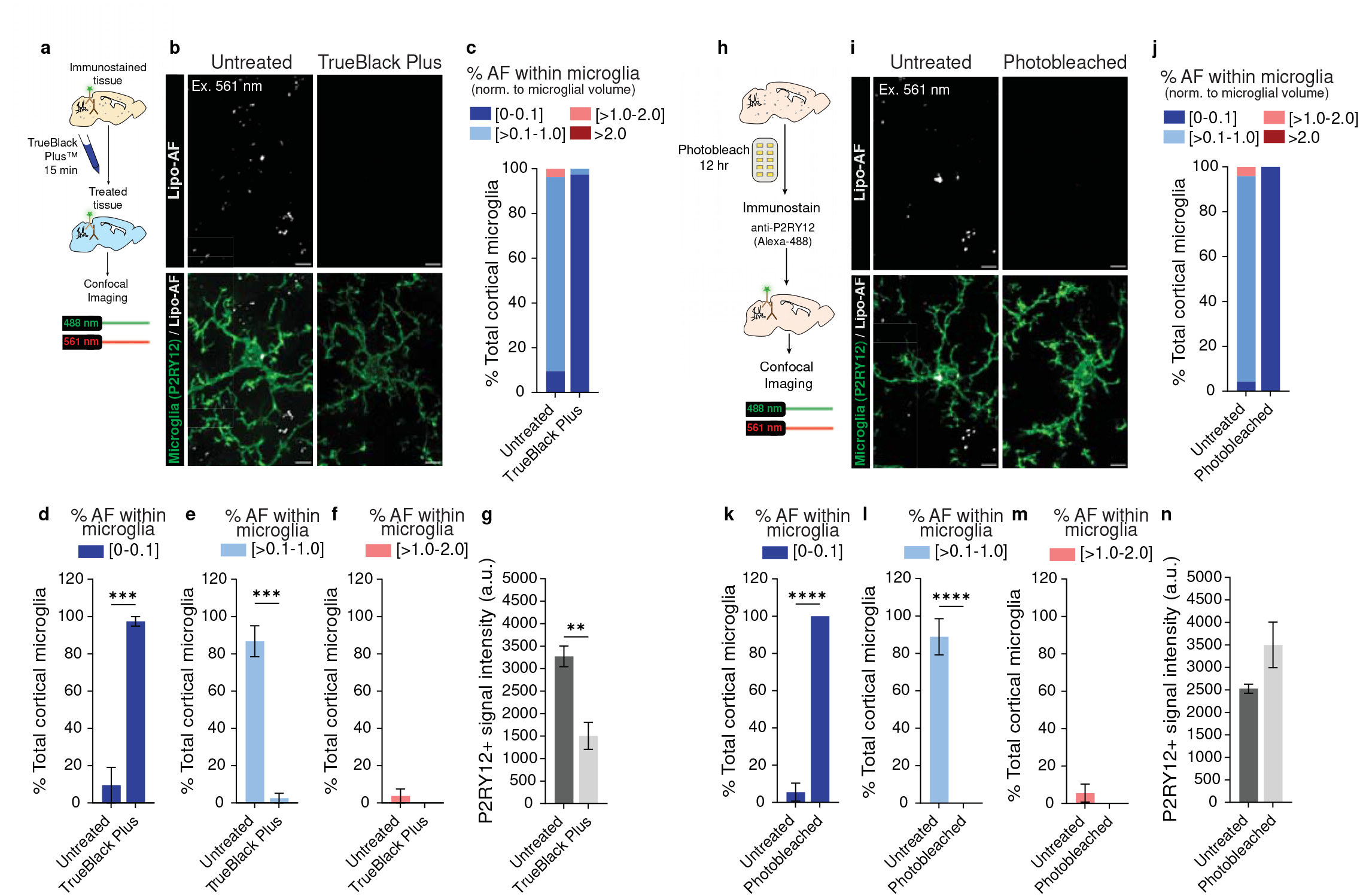
Methods to eliminate lipo-AF signal from mouse brain tissue. **a** Methods for the post-immunostaining quenching protocol using the commercially available TrueBlack Plus^™^ (TBP) reagent. **b** Representative images of anti-P2RY12 immunolabelled microglia (green) and lipo-AF (white) in untreated or TBP-treated samples from P90 mouse cortex. Scale bars = 5 μm. **c** Quantification of the percentage of cortical microglial with [0-0.1]% (dark blue), [>0.1-1.0]% (light blue), [>1.0-2.0]% (pink), or >2.0% (dark red) of their total volume occupied by AF. Data are represented as the mean. **d-f** Same data as **c**, but each bin is graphed separately. Data are represented as a mean ± SEM, n = 3 mice. Two-tailed unpaired t-test (**d**, *t* = 8.912, *df* = 4; **e**, *t* = 9.679, *df* = 4; **f**, *t* = 0.9982, *df* = 4); ***p<0.001. **g** Quantification of the intensity of the anti-P2RY12+ signal in untreated (dark gray bar) or TBP-treated (light gray bar) samples. Data are represented as a mean ± SEM, n = 3 mice. Two-tailed unpaired t-test (*t* = 4.698, *df* = 4); **p<0.01. **h** Schematic describing the pre-immunostaining photobleaching protocol. **I** Representative images of anti-P2RY12 immunolabelled microglia (green) and lipo-AF (white) in untreated or photobleached samples from P90 mouse cortex. Scale bars = 5 μm. **j** Quantification of the percentage of cortical microglial with [0-0.1]% (dark blue), [>0.1-1.0]% (light blue), [>1.0-2.0]% (pink), or >2.0% (dark red) of their total volume occupied by AF. Data are represented as a mean, n = 3 mice. **k-m** Same data as **j**, but each bin is graphed separately. Data represented as a mean ± SEM. Two-tailed unpaired t-test (**k**, *t* = 19.63, *df* = 6; **l**, *t* = 9.235, *df* = 6; **m**, *t* = 1.156, *df* = 6); ****p<0.0001. **n** Quantification of the intensity of the anti-P2RY12+ signal in untreated (dark gray bar) or photobleached (light gray bar) samples. Data presented as mean ± SEM, n = 4 mice. Two-tailed unpaired t-test (*t* = 1.886, *df* = 6).

### Photobleaching can be used to eliminate lipo-AF signal prior to microglial engulfment analyses

Given that lipo-AF is localized within microglial lysosomes (Supplementary Fig. 1), it is possible that it could confound assessments of engulfed cellular material within microglia. As synapses are key phagocytic substrates for microglia in health and disease^1–4^, we used our photobleaching protocol to next determine the impact of lipo-AF on microglial synapse engulfment analysis. Beginning with P90 mouse brain, sections were immunostained for anti-IBA1 to label microglia and anti-vesicular glutamate transporter 2 (VGluT2) to label excitatory presynaptic terminals in layer IV of the somatosensory cortex (Fig. 4a). Neighboring sections from the same brain were left unstained to measure lipo-AF in the same region. A 561nm laser (BP 629/62) was used to image lipo-AF and anti-VGluT2 signal. The intensity of the anti-VGluT2 immunolabelled puncta outside and inside the microglial boundaries, as well as lipo-AF in the neighboring sections, were measured in resulting images (Fig. 4b, immunostained sections are shown). Notably, at P90, the anti-VGluT2 puncta intensity inside the microglia (Fig. 4b,c orange bar) was not significantly different from the lipo-AF intensity captured with the same settings and the same laser line (Fig. 4c gray bar). This suggests that, with the 561 nm laser line (BP 629/62), lipo-AF can yield a comparable signal to anti-VGluT2 immunostaining within microglia and raises the possibility that this can confound analysis of engulfed material within microglia.

**Figure 4.**
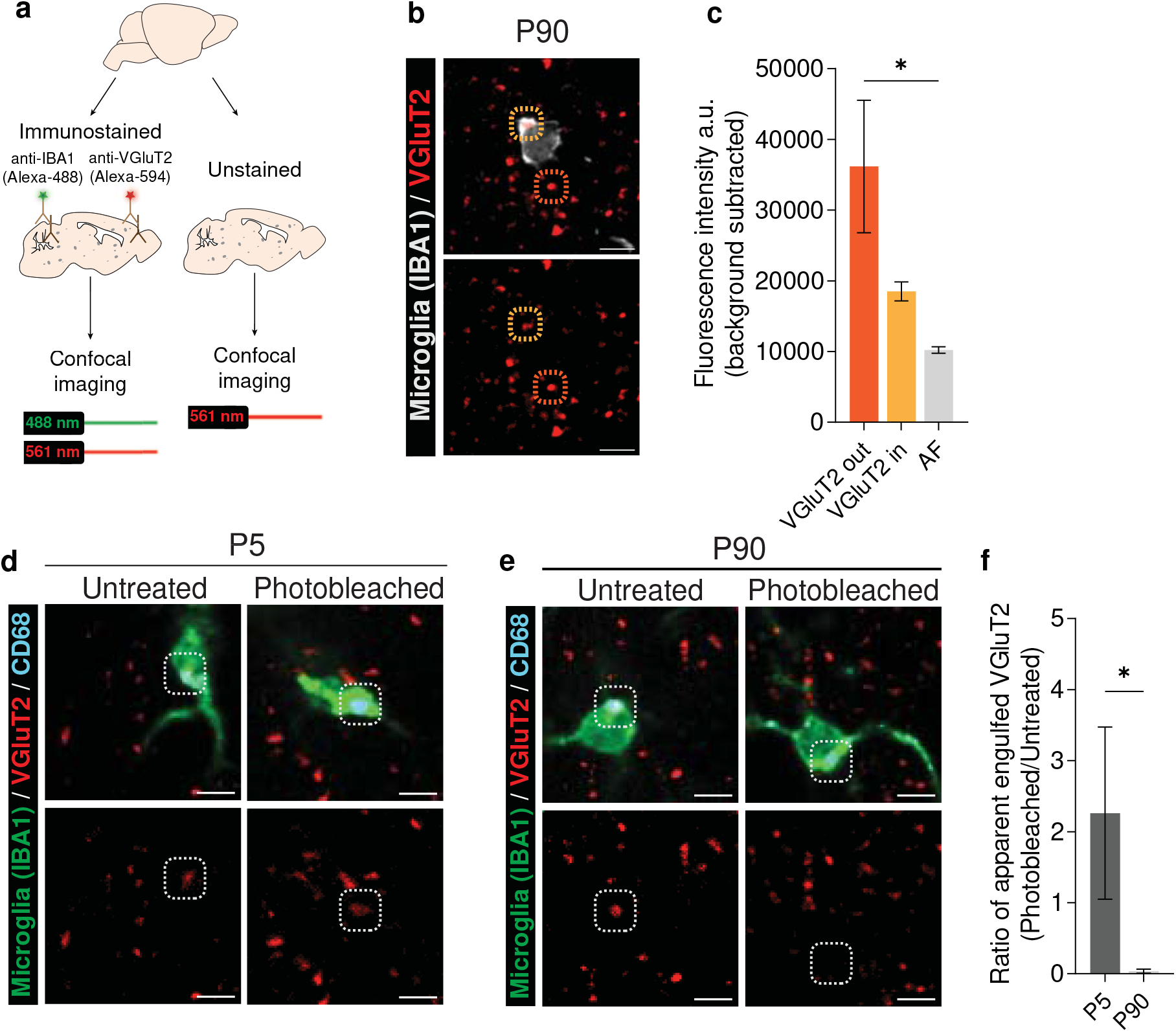
Use of the photobleaching method to determine the degree to which lipo-AF signal can confound analysis of engulfed synaptic material within microglia. **a** P90 mouse brain sections were immunolabeled with anti-VGluT2 antibody and secondary antibodies conjugated with Alexa-Fluor 594. Microglia were also co-immunolabeled with anti-IBA1. Adjacent sections were left unstained to image AF. **b** Representative images of anti-VGluT2+ presynaptic terminals within the P90 somatosensory cortex immunolabelled with Alexa-Fluor 594 secondary antibody. The dashed orange circles highlight the anti-VGluT2 signal outside microglia. The dashed yellow circles highlight the anti-VGluT2 signal inside microglia. Bottom image is the same image without the anti-IBA1 immunostaining. Scale bars = 5 μm. **c** Quantification of the fluorescence intensity of Alexa-Fluor 594 immunolabelled VGluT2+ puncta, outside (orange bar) and inside microglia (yellow bar) upon excitation with a 561 nm laser line. For comparison, lipo-AF signal intensity (gray bar), upon excitation with a 561 nm laser line, in adjacent tissue sections was also quantified. Data are represented as a mean ± SEM, n = 3 mice. One-way ANOVA with Tukey multiple comparisons test (*F* = 5.873, *df* = 8); *p<0.05. **d-e** Representative images of signal in the anti-VGluT2+ immunolabeled channel (red) inside anti-CD68+ lysosomes (cyan) of anti-IBA1 immunolabelled microglia (green) in untreated (left) or photobleached samples (right) from P5 (d) or P90 (e) mouse cortex. White dotted circles highlight signal within the anti-VGluT2 channel that co-localize with microglial C68+ lysosomes. Bottom images in d and e are the same images as the top images without the anti-IBA1 immunostaining. Scale bars = 5 μm. **f** The ratio of the volume of apparent engulfed anti-VGluT2+ material within microglia from the P5 and P90 mouse cortex post-photobleaching to pre-photobleaching (volume of anti-VGluT2 signal within microglia post-photobleaching/pre-photobleaching signal). Data are represented as a mean ± SEM, n = 4 mice. Two-tailed Mann-Whitney t-test; *p<0.05.

We more directly tested the extent to which the anti-VGluT2 signal detected within the microglial boundaries could be confounded by lipo-AF signal using the photobleaching protocol (Fig. 3h). We first photobleached P90 tissue to rid of AF and then immunostained tissue for VGluT2 and IBA1. We imaged all tissues with a 561nm laser (BP 629/62) in the P90 and P5 mouse somatosensory cortex. We chose to compare P90 to P5 as P5 was a developmental timepoint where lipo-AF was low (Fig. 1), and it is an age where the somatosensory cortex is known to undergo extensive experience-dependent synapse remodeling by phagocytic microglia^25^. In untreated sections, apparent engulfed VGluT2 material was detectable within microglia at P5 and P90 (Fig. 4d-e). However, after photobleaching, this engulfed VGluT2 signal was no longer detected at P90 (Fig. 4d,f). In contrast, microglia within the P5 cortex displayed similar levels of engulfed VGluT2+ material within their cytosol in the photobleached and non-photobleached condition (Fig. 4e-f). We further compared the fold difference in signal intensity of anti-VGluT2 signal within microglia after photobleaching to without photobleaching (post-photobleaching anti-VGluT2+ signal within microglia/pre-photobleaching signal). There was a significant reduction in anti-VGluT2 within microglia in the P90 cortex after photobleaching compared to P5 (Fig. 4f). Together, these data suggest that lipo-AF can confound the interpretation of fluorescent signal within microglial lysosomes in young adult mouse brain tissue, but this is less of a concern in neonate brain tissue. Nonetheless, precautions should be taken to eliminate lipo-AF to avoid false positive detection of engulfment events.

### Photobleaching eliminates autofluorescence in aged mouse, marmoset, and human brain tissue

Photobleaching effectively quenched lipo-AF in mouse tissue (Figs. 3–4). We, thus, extended this protocol to older mouse tissue and to other species (Fig. 5). We found that extending the photobleaching period to 24 hours significantly reduced AF signal across multiple fluorescence channels in 24-month-old aged mouse cortex (Fig. 5a,e) and 9-month-old 5xFAD mouse cortex (Fig. 5b, f). Similar to our experiments in mice, we found that photobleaching for 24 hours significantly reduced the signal intensity from AF across multiple fluorescence channels in formalin-fixed, paraffin-embedded (FFPE) 11–13 year-old marmoset cortex (Fig. 5c,g) and FFPE 60–77 year-old human cortex (Fig. 5d,h). Note, the human tissue was collected from the postmortem brains of multiple sclerosis (MS) subjects. Together, we have implemented a new pre-staining protocol that reliably eliminates AF in tissue sections in different tissue preparations. As this protocol can be adapted for multiple species tissues, including human, the protocol has broad applicability.

**Fig 5.**
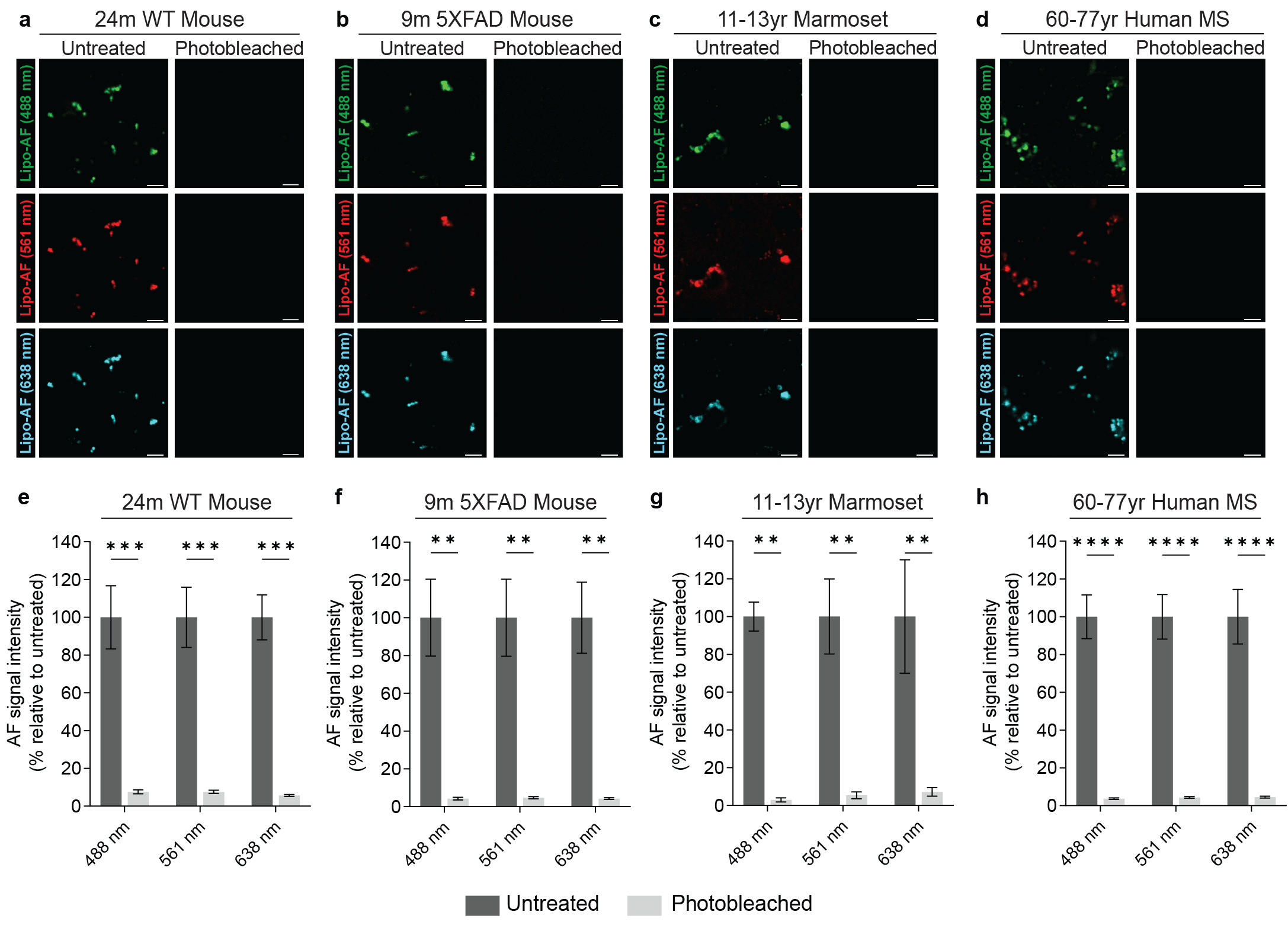
Photobleaching quenches lipo-AF signal in mouse, marmoset and human brain tissue. **a-d** Representative images of lipo-AF upon excitation with a 488 nm, 561 nm or 638 nm laser line in untreated (left column) or photobleached (right column) samples from 24-month-old mouse cortex (a), 9-month-old 5XFAD mouse cortex (b), 11-13 year-old marmoset cortex (c), and 60-77 year-old human cortex (d). Scale bars = 5 μm. **e-h** Quantification of lipo-AF signal intensity before and after photobleaching for each species. Data represented as a mean ± SEM, n = 3 biological replicates per species. Two-way ANOVA with Šídák’s multiple comparisons test (**e**, *F* = 114.7, *df* = 1; **d**, *F* = 69.41, *df* = 1; **f**, *F* = 59.30, *df* = 1; **h**, *F* = 171.3, *df* = 1;); **p<0.01, ***p<0.001****p<0.0001.

## Discussion

Here, we assessed microglia-associated lipo-AF in the developing, adult, aged, and diseased mouse cortex using confocal light microscopy. A small degree of microglial lipo-AF could be detected as early as P5 and microglia were the first cells in the mouse cortex to accumulate lipo-AF. Lipo-AF within microglia increased into adulthood and a large accumulation of lipo-AF was detected inside and outside of microglia in the aged mouse cortex. We further showed that lipo-AF within microglia can be mis-interpreted as engulfed synaptic material, particularly in the young adult mouse brain. Finally, we provide a new protocol to rid tissues of AF signal before immunostaining to reduce the confound of lipo-AF for microglial engulfment studies, which can also be applied to any other study reliant on fluorescence light microscopy. Importantly, we show that this AF quenching protocol is broadly applicable and can be performed in mouse, non-human primate, and human brain tissues.

One surprising result from this study is that neurodegeneration and Aβ accumulation in 5XFAD mice did not exacerbate the accumulation of lipo-AF in microglia. One study has found that, following microglia depletion, the proliferating microglia were primarily restricted to the AF-negative, likely lipofuscin-negative, subset of cells^17^. It is also known that cell division is the only way cells can reduce lipofuscin, which is likely to underlie the lipo-AF signal we are observing in brain tissue^13,14,22^. That is, as cells divide, lipofuscin is diluted. Thus, it is possible that lipofuscin-negative microglia or microglia with low amounts of lipofuscin proliferate during neurodegeneration and, thus, the amount of lipofuscin and AF due to lipofuscin per cell is decreased. Indeed, it has been previously shown in 5XFAD mice that cortical microglia increase, and microglia near Aβ plaques proliferate^26,27^. It is also possible that lipo-AF accumulation is disease and/or substrate specific. For example, the engulfment of myelin in a demyelinating disease related to MS has been suggested to drive lipofuscin accumulation in microglia^19^. Therefore, specific engulfed substrates may lead to the accumulation of lipofuscin, while others do not. An important future direction will be to understand the prevalence of this biology in microglia across different diseases.

What is the source of lipo-AF, and how does it accumulate in microglia? We speculate that the broad phagocytic role of microglia in early development may explain the presence of the small amount of lipo-AF within microglia in the early postnatal cortex. This lipo-AF is likely from lipofuscin given its excitation and emission spectra. It is possible that the lysosomal degradation capacity of microglia in the young brain may be able to compensate and/or microglia are actively dividing in the young brain and, thereby, lipofuscin is being diluted. In contrast, during adulthood, ongoing engulfment of cellular material and decreased cell division may lead to accumulation of lipofuscin and its related AF. This may also increase with aging as a result of age-related lysosomal dysfunction^12^. Indeed, recent studies have shown how disruption of the lysosomal degradation pathway in microglia during engulfment can contribute to lipofuscin accumulation^19^. Better defining the molecular composition and accumulation of these lipo-AF deposits in microglia throughout the lifespan and disease would be most informative.^17, 19^

Another important aspect of our study is that we provide two distinct methods to eliminate AF signal from tissue sections before imaging. The first uses the commercially available reagent TrueBlack Plus^™^, an alternative to Sudan Black dye. Our data shows that a 15-minute post-staining treatment was sufficient to eliminate the AF signal. However, we also found a significant quenching of the immunostained fluorophore signal. The second method consisted of incubating sections in a commercially available device that administers light to a sample for an extended period. Using this device, we found that photobleaching brain tissue sections with light for 12-24 hours significantly reduces the AF signal without compromising fluorophore signal intensity. While our device’s intensity and wavelength of light remains proprietary, previous studies have used a similar photobleaching strategy with LEDs^24, 21,28^. Thus, this technology could be easily adapted across laboratories and tissue samples. Finally, we applied the photobleaching protocol in the context of microglial synapse engulfment analysis^10^. Several studies have shown how microglia engulf synaptic material at neonatal time points^1,3,4^. Despite detecting low levels of microglial lipo-AF in the early postnatal P5 brain, it had no confounding effect on the analysis of VGluT2 engulfment within microglia. That is, anti-VGluT2 immunofluorescence signal was still observed within microglia after photobleaching samples. Conversely, in young, adult mice (P90), apparent engulfed VGluT2 within microglia was no longer observed after photobleaching. Therefore, caution should be used when interpreting microglial engulfment, particularly in adult mouse brain. It is noted that, although we detected lipo-AF with 3 different laser lines (Fig. 1), we did most of our analyses with the 561nm laser (BP 629/62). There may be variations in detecting lipo-AF on other microscope set ups and with other laser lines and filter sets. Still, photobleaching samples before immunostaining and including secondary-only controls to evaluate the amount of lipofuscin-derived AF or other sources of AF in tissue sections are the best practice. This is particularly important considering that engulfed material and lipo-AF share the same subcellular compartment (i.e., lysosomes) within microglia.

In summary, as more and more studies are realizing the impact of microglial engulfment mechanisms on neural circuit structure and function^1–4^, it is critical to perform experiments to assess engulfment of cellular and protein substrates by microglia to the highest rigor. The protocols we provide ensure that microglial engulfment confocal imaging assays are not confounded by AF. Importantly, these protocols can be used in mouse models, but the protocols can also be adapted for use in non-human primate and human tissue samples.

## Materials and Methods

### Animals

Male and female wildtype C57Bl/6J mice (stock #000664) were obtained from Jackson Laboratories (Bar Harbor, ME). Adult common marmosets (*Callithrix jacchus*), both male and females between 11-13 years old, were obtained from the marmoset tissue library of translational neuroradiology section (TNS) at the NINDS. All animal experiments were performed in accordance with Animal Care and Use Committees (IACUC) and under NIH guidelines for proper animal welfare.

### Human samples

Collection of human multiple sclerosis (MS) postmortem brain tissue was performed after obtaining informed consent for collection and were obtained from the Translational Neuroradiology Section at the NIH/NINDS. Samples analyzed in the current study were collected from the insular/parietal neocortex and prefrontal cortex of 3 women with multiple sclerosis (MS) with ages ranging from 60–77 years.

### Immunostaining

Mice were anesthetized and transcardially perfused with 0.1M phosphate buffer (PB) followed by 4% paraformaldehyde (PFA) (Electron Microscopy Services 15710)/0.1M PB. Brains were post-fixed at 4°C in PFA overnight, equilibrated in 30% sucrose/0.1M PB and then embedded in a 2:1 mixture of 30% sucrose/0.1M PB and O.C.T. compound (ThermoFisher Scientific Waltham, MA, USA). To ensure methods were of global use, sections were immunostained on slides or floating. A cryostat was used to cut either 10-16 μm coronal brain sections on slides (microglial lipo-AF analysis across development and in 5XFAD mice) or 40 μm floating sections in 0.1M PB (lipo-AF intensity, quenching and synaptic engulfment analysis). Subsequent sections were blocked and permeabilized at room temperature for 1 hr in blocking solution (10% normal goat serum/0.1M PB containing 0.3% Triton-X 100) followed by overnight incubation with primary antibodies at ambient room temperature. Primary antibodies included: Rat mAb anti-CD68 (Abcam, ab955; 1:200), rabbit pAb anti-IBA1 (Wako Chemicals, 019-19741; 1:500), chicken mAb anti-IBA1 (Synaptic Systems, 234009; 1:500), rabbit pAb anti-P2RY12 (Anaspec, 55043A; 1:2000) and guinea pig pAb anti-VGluT2 (Millipore, Ab2251-I; 1:1000). The following day, sections were washed 3×5 min with 0.1M PB and incubated with the appropriate Alexa-fluorophore-conjugated secondary antibodies including goat anti-chicken IgY (H+L) Alexa-Fluor 488 IgY (Life Technologies Scientific; A11039), goat anti-rabbit IgG (H+L) Alexa-Fluor 488 (Life Technologies; A11034), goat anti-guinea pig IgG (H+L) Alexa-Fluor 488 (Life Technologies; A11073), goat anti-rabbit IgG (H+L) Alexa-Fluor 594 (Life Technologies; A11012), goat anti-guinea pig IgG (H+L) Alexa-Fluor 594 (Life Technologies; A11076), goat anti-rabbit IgG (H+L) Alexa-Fluor 647 (Life Technologies; A21245), goat anti-guinea pig IgG (H+L) Alexa-Fluor 647 (Life Technologies; A21450), goat anti-rat IgG (H+L) Alexa-Fluor 647 (Life Technologies; A21247) for 2 hr at room temperature. Slides and floating sections were washed 3×10 min with 0.1M PB. Floating sections were then mounted on slides. All subsequent slides were air dried and cover glass (ThermoFisher; 12-544-DP) was mounted with Vectashield containing DAPI (Vector laboratories, Burlingame, CA, USA) or with CFM-3 (Citifluor, Hatfield, PA, USA) for chemical quenching experiments.

### Chemical quenching

After staining, floating sections or tissue-mounted slides were incubated with TrueBlack^®^ Plus Lipofuscin Autofluorescence Quencher (Biotium, Fremont, CA, USA) in 1X phosphate buffer saline (PBS) for 15 min at room temperature with rocking. Following 3×5 min washes with PBS 1X, sections were mounted with CFM-3 (Citifluor, Hatfield, PA, USA).

### Photobleaching

Before incubating in blocking solution, floating sections or tissue-mounted slides were placed in 0.1M PB and incubated in the MERSCOPE Photobleacher (Vizgen, Cambridge, MA, USA) for 12-24 hour. Photobleached samples were then incubated in blocking solution and immunostained as described above.

### Confocal Imaging

Mounted brain sections were imaged using Zen Blue acquisition software (Zeiss; Oberkochen, Germany) on a Zeiss Observer Spinning Disk confocal microscope equipped with diode lasers 405 nm/50 mW, 488 nm/50 mW, 561 nm/50 mW, and 638 nm/75 mW, and with 450/50 (blue), 525/50 (green), 629/62 (red) and 690/50 (far-red) BP emission filter sets, respectively. For most experiments, 6-12 (AF) or 3 (anti-VGluT2 immunostained sections) 40x fields of view were randomly chosen within the somatosensory and neighboring visual and auditory cortices an z-stacks were acquired at 0.31 μm spacing. For AF, anti-VGluT2, and anti-P2RY12 intensity measurements, 2-4 63x fields of view were randomly and z-stacks were then acquired at 0.27 μm spacing. For all imaging experiments, identical settings were used to acquire images from all samples within one experiment.

### Quantification of lipo-AF and VGluT2 volume

Images were first pre-processed and blinded using a custom macro in ImageJ (NIH, version 1.53c). Imaris v9 (Bitplane) was then used to create a 3D surface rendering of single microglia cells. A 3D surface rendering of masked anti-CD68+ signal from each cell was also created and was used to generate a 3D surface rendering of CD68-masked AF+ or CD68-masked VGluT2+ material. Lipo-AF or VGluT2 within microglia was calculated by dividing the volume of CD68-masked lipo-AF or VGluT2 by the volume of the individual microglial cell. To quantify lipo-AF or VGluT2 signal outside the microglia volume, a 3D surface rendering of total lipo-AF or VGlut2 was created and the volume of lipo-AF or VGluT2 within microglia was then subtracted.

### Quantification of VGluT2 and lipo-AF signal intensity

All analyses of intensity were performed on single z-planes blind to condition using ImageJ (NIH, version 1.53c). Briefly, for sections stained with anti-VGluT2 and anti-IBA1, square regions of interests (ROIs) of equal size were drawn at the center of anti-IBA1 immunolabelled microglial somas containing anti-VGluT2 positive signal. Within these ROIs, smaller circular ROIs of equal size were placed at the center of anti-VGluT2 immunolabelled puncta outside and inside anti-IBA1 immunolabelled microglia. Three background circular ROIs of the same size were also selected for each field of view within the single z-plane. The raw integrated density of pixels within each circular ROI was measured. To quantify lipo-AF intensity, unstained sections were imaged and subsequently ROIs of equal size were drawn at the center of single z-planes containing multiple lipo-AF puncta. Similar to immunostained sections, circular ROIs of equal size were then placed at the center of AF. Three background ROIs were also selected. For all images (immunostained with VGluT2 or unstained to measure AF), the raw integrated density of pixels within each circular ROI was measured and the background ROI pixel intensity value was averaged. Last, for each VGluT+ or AF+ puncta intensity measurement, the average background was subtracted prior to statistical comparison.

### Quantification of P2Y12R signal intensity

To quantify the fluorescence intensity of anti-P2Y12R signal, Imaris v9 (Bitplane) was used to create a 3D surface rendering of all microglia cells and the intensity mean parameter of anti-P2Y12R signal was then extracted.

### Quantification of AF signal intensity after photobleaching in mouse, marmoset, and human brain tissue

Unstained sections were imaged on a Zeiss Observer Spinning Disk confocal microscope as described above. All subsequent analyses of intensity were performed on single z-planes using ImageJ (NIH, version 1.53c). For each image, the 488 nm laser channel was background subtracted (10x) and despeckled (median filter, 3×3) in ImageJ and was then ROIs were selected using the analyze particle function. These ROIs were then applied across all channels for that image. The raw integrated density of pixels within each individual ROI was then measured for each fluorescence channel. The average raw integrated density of pixels for each of three z-stacks was then calculated, and the average per biological replicate and condition were calculated.

### Statistical analysis

Results are presented as either mean or mean ± standard error (SEM). For normally distributed data, analyses included two-tailed, unpaired Students t-test when comparing 2 conditions or one-way ANOVA followed by Tukey’s post hoc analysis or two-way ANOVA followed by Sidak’s post hoc analyses (indicated in figure legends) using GraphPad Prism (La Jolla, CA). For data that were not normally distributed, a Mann-Whitney test was used. Values of *p < 0.05, **P < 0.01, ***P < 0.001, ****P < 0.0001 were considered statistically significant.

## Acknowledgments

We thank Dr. Christina Baer (UMMS) for critical reading of the manuscript and assistance with microscopy. We thank Joan Ohayon, CRNP, for coordinating MS autopsies. This work was supported by NIMH-R01MH113743 (DPS), NINDS-R01NS117533 (DPS), NIA-RF1AG068281 (DPS), NIH-T32AI132152 (JMS), the Intramural Research Program of NINDS (DSR), and the Dr. Miriam and Sheldon G. Adelson Medical Research Foundation (DPS and DSR).

## Author contributions

J.M.S., F.M.L, and D.P.S. designed the study and wrote the manuscript. J.M.S., F.M.L, and D.P.S. performed and analyzed the experiments. D.S.R. supervised collection and processing of marmoset and human brain tissue by J.L. and K.H. All authors edited and revised the manuscript.

## Materials & Correspondence

All materials and correspondences should be addressed to Dr. Dorothy P. Schafer.

## References

1 Schafer, D. P. & Stevens, B. Phagocytic glial cells: sculpting synaptic circuits in the developing nervous system. Current opinion in neurobiology 23, 1034–1040, doi:10.1016/j.conb.2013.09.012 (2013).

2 Hong, S., Dissing-Olesen, L. & Stevens, B. New insights on the role of microglia in synaptic pruning in health and disease. Current opinion in neurobiology 36, 128–134, doi:10.1016/j.conb.2015.12.004 (2016).

3 Stevens, B. & Schafer, D. P. Roles of microglia in nervous system development, plasticity, and disease. Dev Neurobiol 78, 559–560, doi:10.1002/dneu.22594 (2018).

4 Faust, T. E., Gunner, G. & Schafer, D. P. Mechanisms governing activity-dependent synaptic pruning in the developing mammalian CNS. Nature reviews. Neuroscience 22, 657–673, doi:10.1038/s41583-021-00507-y (2021).

5 Crapser, J. D., Arreola, M. A., Tsourmas, K. I. & Green, K. N. Microglia as hackers of the matrix: sculpting synapses and the extracellular space. Cell Mol Immunol 18, 2472–2488, doi:10.1038/s41423-021-00751-3 (2021).

6 Song, W. M. & Colonna, M. The identity and function of microglia in neurodegeneration. Nat Immunol 19, 1048–1058, doi:10.1038/s41590-018-0212-1 (2018).

7 Germann, M., Brederoo, S. G. & Sommer, I. E. C. Abnormal synaptic pruning during adolescence underlying the development of psychotic disorders. Curr Opin Psychiatry 34, 222–227, doi:10.1097/YCO.0000000000000696 (2021).

8 Biber, K., Moller, T., Boddeke, E. & Prinz, M. Central nervous system myeloid cells as drug targets: current status and translational challenges. Nature reviews. Drug discovery 15, 110–124, doi:10.1038/nrd.2015.14 (2016).

9 Priller, J. & Prinz, M. Targeting microglia in brain disorders. Science 365, 32–33, doi:10.1126/science.aau9100 (2019).

10 Schafer, D. P., Lehrman, E. K., Heller, C. T. & Stevens, B. An engulfment assay: a protocol to assess interactions between CNS phagocytes and neurons. Journal of visualized experiments: JoVE, doi:10.3791/51482 (2014).

11 Morini, R., Bizzotto, M., Perrucci, F., Filipello, F. & Matteoli, M. Strategies and Tools for Studying Microglial-Mediated Synapse Elimination and Refinement. Front Immunol 12, 640937, doi:10.3389/fimmu.2021.640937 (2021).

12 Moreno-Garcia, A., Kun, A., Calero, O., Medina, M. & Calero, M. An Overview of the Role of Lipofuscin in Age-Related Neurodegeneration. Front Neurosci 12, 464, doi:10.3389/fnins.2018.00464 (2018).

13 Brunk, U. T. & Terman, A. Lipofuscin: mechanisms of age-related accumulation and influence on cell function. Free Radic Biol Med 33, 611–619, doi:10.1016/s0891-5849(02)00959-0 (2002).

14 Gray, D. A. & Woulfe, J. Lipofuscin and aging: a matter of toxic waste. Sci Aging Knowledge Environ 2005, re1, doi:10.1126/sageke.2005.5.re1 (2005).

15 Calin, E. F. et al. Lipofuscin: a key compound in ophthalmic practice. Rom J Ophthalmol 65, 109–113, doi:10.22336/rjo.2021.23 (2021).

16 Mochizuki, Y., Park, M. K., Mori, T. & Kawashima, S. The difference in autofluorescence features of lipofuscin between brain and adrenal. Zoolog Sci 12, 283–288, doi:10.2108/zsj.12.283 (1995).

17 Burns, J. C. et al. Differential accumulation of storage bodies with aging defines discrete subsets of microglia in the healthy brain. Elife 9, doi:10.7554/eLife.57495 (2020).

18 Zhang, H., Tan, C., Shi, X. & Xu, J. Impacts of autofluorescence on fluorescence based techniques to study microglia. BMC Neurosci 23, 21, doi:10.1186/s12868-022-00703-1 (2022).

19 Safaiyan, S. et al. Age-related myelin degradation burdens the clearance function of microglia during aging. Nat Neurosci 19, 995–998, doi:10.1038/nn.4325 (2016).

20 Sierra, A., Gottfried-Blackmore, A. C., McEwen, B. S. & Bulloch, K. Microglia derived from aging mice exhibit an altered inflammatory profile. Glia 55, 412–424, doi:10.1002/glia.20468 (2007).

21 Tsuneoka, Y., Atsumi, Y., Makanae, A., Yashiro, M. & Funato, H. Fluorescence quenching by high-power LEDs for highly sensitive fluorescence in situ hybridization. Frontiers in molecular neuroscience 15, 976349, doi:10.3389/fnmol.2022.976349 (2022).

22 Terman, A. & Brunk, U. T. Lipofuscin: mechanisms of formation and increase with age. APMIS 106, 265–276, doi:10.1111/j.1699-0463.1998.tb01346.x (1998).

23 Croce, A. C. & Bottiroli, G. Autofluorescence spectroscopy and imaging: a tool for biomedical research and diagnosis. Eur J Histochem 58, 2461, doi:10.4081/ejh.2014.2461 (2014).

24 Duong, H. & Han, M. A multispectral LED array for the reduction of background autofluorescence in brain tissue. J Neurosci Methods 220, 46–54, doi:10.1016/j.jneumeth.2013.08.018 (2013).

25 Gunner, G. et al. Sensory lesioning induces microglial synapse elimination via ADAM10 and fractalkine signaling. Nature neuroscience 22, 1075–1088, doi:10.1038/s41593-019-0419-y (2019).

26 Forner, S. et al. Systematic phenotyping and characterization of the 5xFAD mouse model of Alzheimer’s disease. Sci Data 8, 270, doi:10.1038/s41597-021-01054-y (2021).

27 Wang, Y. et al. TREM2-mediated early microglial response limits diffusion and toxicity of amyloid plaques. The Journal of experimental medicine 213, 667–675, doi:10.1084/jem.20151948 (2016).

28 Sun, Y.’ Ip, P. & Chakrabartty, A. Simple Elimination of Background Fluorescence in Formalin-Fixed Human Brain Tissue for Immunofluorescence Microscopy. J Vis Exp, doi:10.3791/56188 (2017).

