## Supplementary Figure 1 for "Lipofuscin-like autofluorescence within microglia and its impact on studying microglial engulfment"

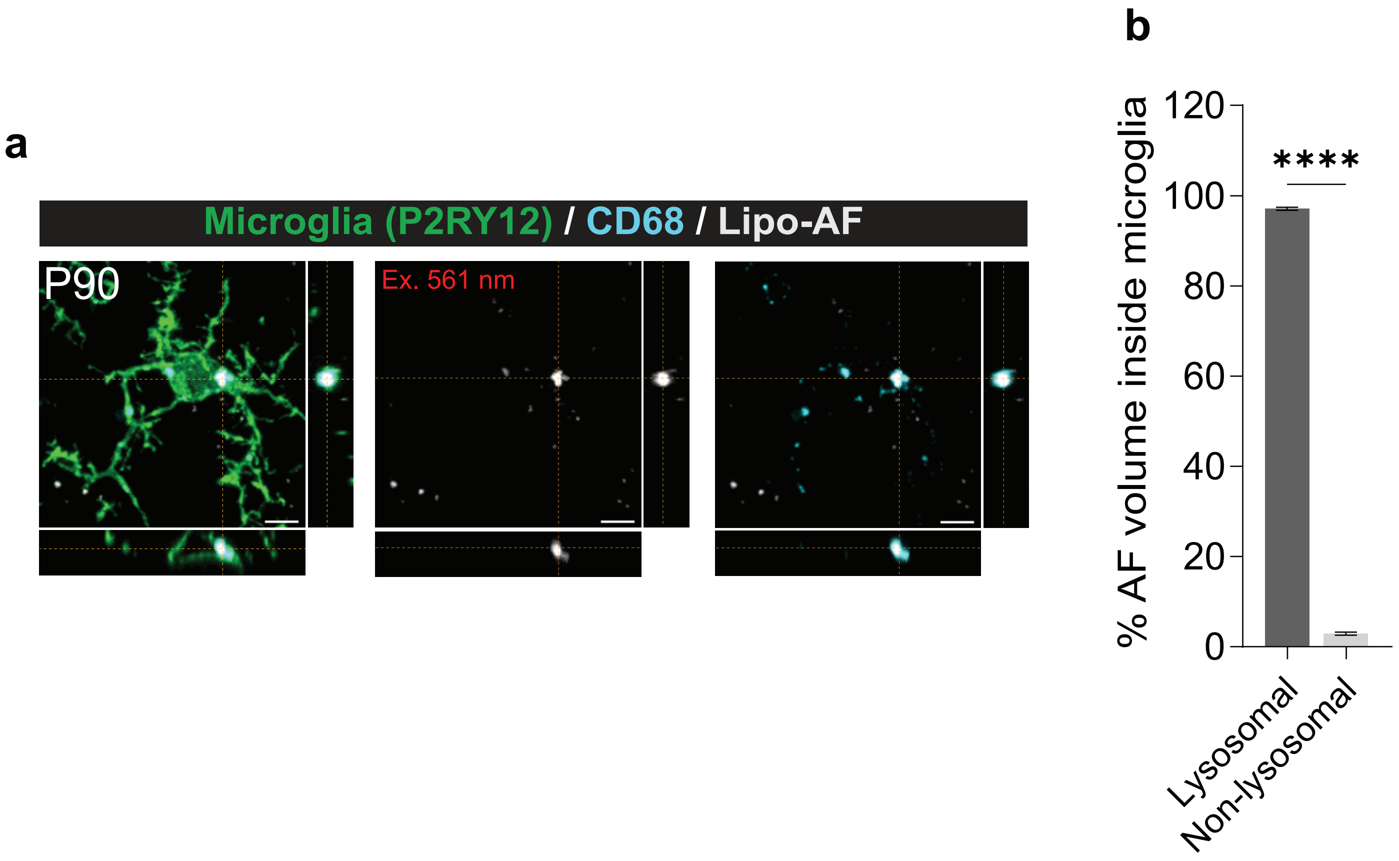
a
Microglia (P2RY12) / CD68 / Lipo-AF
P90
Ex. 561 nm
b
% AF volume inside microglia
120
100
80
60
40
20
0
****
Lysosomal
Non-lysosomal

**Supplementary Figure 1. Microglial lipo-AF is enriched in CD68+ lysosomes. a** Representative orthogonal view of an anti-P2RY12 immunolabelled microglia (green) in the P90 mouse cortex containing lipofuscin-like autofluorescence (lipo-AF, white) within anti-CD68+ lysosomal compartments (cyan). Scale bars = 5 μm. **b** Quantification of the percentage of lysosomal (dark gray bar) and non-lysosomal (light gray bar) lipo-AF inside microglia. Data are represented as mean ± SEM, n = 3 mice. Two-tailed unpaired t-test ($t$ = 198.5, $df$ = 4); ****$p<0.0001$.
